# A peptide targeting outer membrane protein A of *Acinetobacter baumannii* exhibits antibacterial activity by reducing bacterial pathogenicity

**DOI:** 10.1101/2024.04.03.587999

**Authors:** Hui Zhao, Yue Hu, Dan Nie, Zhou Chen, Na Li, Shan Zhou, Mingkai Li, Xiaoyan Xue

## Abstract

The World Health Organization has classified multidrug-resistant (MDR) *Acinetobacter baumannii* as a significant threat to human health, necessitating the urgent discovery of new antibacterial drugs to combat bacterial resistance. Outer membrane protein A of *A. baumannii* (AbOmpA) is an outer membrane-anchored β-barrel-shaped pore protein that plays a critical role in bacterial adhesion, invasion, and biofilm formation. Therefore, AbOmpA is considered a key virulence factor of *A. baumannii*. Herein, we screened three phage display peptide libraries targeting AbOmpA and identified several peptides. Among them, P92 (amino acid sequence: QMGFMTSPKHSV) exhibited the highest binding affinity with AbOmpA, with a KD value of 7.84 nM. In vitro studies demonstrated that while P92 did not directly inhibit bacterial growth, it significantly reduced the invasion and adhesion capabilities of multiple clinical isolates of MDR *A. baumannii* and concentration-dependently inhibited biofilm formation by acting on OmpA. Furthermore, the polymerase chain reaction results confirmed a significant positive correlation between the antibacterial effect of P92 and OmpA expression levels. Encouragingly, P92 also displayed remarkable therapeutic efficacy against *A. baumannii* infection in various models, including an in vitro cell infection model, a mouse skin infection model, and a mouse sepsis model. These results highlight P92 as a novel and highly effective antimicrobial molecule specifically targeting the virulence factor AbOmpA.

**IMPORTANCE:** Different from the action mechanism of traditional antibiotics, antibacterial strategies aimed at targeting bacterial virulence factors can effectively reduce bacterial pathogenicity without compromising bacterial growth and reproduction, making it a crucial research direction in combating bacterial drug resistance. Despite the influence of various factors on the expression of bacterial virulence factors, specific and consistently expressed virulence factors in certain bacteria can still serve as viable targets for drug development. In this study, we focused on outer membrane protein A, a key virulence factor of *A. baumanni*i, and successfully identified a highly specific antimicrobial peptide P92 through screening processes. Furthermore, our findings demonstrate its efficacy in various dynamic models for anti-infective therapy. These results validate that antibacterial agents targeting bacterial virulence factors possess relatively or absolutely narrow spectrum antimicrobial properties, enabling precise antibacterial action without inducing bacterial resistance.

## Introduction

Multi-drug resistant (MDR) gram-negative bacteria have emerged as a prominent cause of mortality resulting from severe infections. In 2017, the World Health Organization released a list comprising 12 species of antibiotic-resistant bacteria, with 9 belonging to the category of gram-negative bacteria (e.g. *Acinetobacter baumannii*, *Pseudomonas aeruginosa*, and *Enterobacter*). This list highlights the importance of identifying novel targets and developing new drugs against MDR gram-negative bacteria. The outer membrane (OM), which constitutes an atypical asymmetric phospholipid bilayer, serves as a distinctive structure in gram-negative bacteria. The inner leaflet consists of glycerophospholipids, while the outer leaflet comprises lipopolysaccharides. This bilayer acts as a formidable barrier that hinders the diffusion of most drug molecules across the membrane(1, 2). The OM encompasses various proteins known as outer membrane proteins (OMPs) that are essential for bacterial growth and pathogenesis, which include lipopolysaccharide (LPS) and capsular polysaccharide transporters, porins facilitating macromolecular uptake, and transporters responsible for vital elements such as iron(3). OMPs play a critical role in maintaining structural and functional integrity within the OM of gram-negative bacteria(4). The antibiotic murepavadin, which specifically targets lipopolysaccharide transport protein D (LptD), or chimeric peptidomimetics that target BamA (involved in OMP transport and assembly) exhibit remarkable efficacy against MDR gram-negative bacterial strains in vivo and in vitro(1). Therefore, significant research interest is focused on developing novel antibacterial drugs that target specific functions associated with OMP molecules.

In *A. baumannii,* the primary OM porin is monomeric OmpA, which functions as a nonspecific slow porin and plays a critical role in regulating the transmembrane transport of small molecule nutrients and hydrophilic antimicrobial drugs(5). Moreover, it serves as a pivotal virulence factor for *A. baumannii*, being closely associated with bacterial adhesion and invasion, biofilm formation, immune evasion, and survival in harsh environments(4, 6). Furthermore, OmpA contributes to bacterial resistance; studies have reported that the knockout of the *ompA* gene leads to an 8-fold reduction in the minimum inhibitory concentration (MIC) of aztreonam against MDR *A. baumannii*(7). It is noteworthy that that inhibition of OmpA represents an antibacterial virulence strategy that is distinct from traditional antibiotic modes of action. Antivirulence molecules do not produce direct bactericidal effects or survival pressure on bacteria, and therefore do not induce bacterial resistance.

Currently, various antibacterial strategies targeting OmpA have been reported, encompassing peptides, vaccines, and antibodies(8). For example, a cyclic hexapeptide, AOA-2(9) has not only demonstrated the ability to reduce the adhesion of *A. baumannii*, *P. aeruginosa*, and *E. coli* to the surfaces of biotic and abiotic agents but also exhibited synergistic antibacterial activity with colistin. Herein, we used the recombinant *A. baumannii* OmpA (AbOmpA) as the target, and identified six peptides by screening from the phage displayed random peptide libraries. Through comprehensive in vivo and in vitro investigations, we demonstrated that peptide-P92 exhibited the highest affinity for AbOmpA thus far and effectively inhibited virulence in MDR strains of *A. baumannii*. Our results strongly support P92 as a novel and highly effective antimicrobial molecule with promising prospects for anti-infection therapy.

## Results

### Screening and evaluation of antiAbOmpA peptides

First, we constructed a pET30a(+)/AbOmpA plasmid. Then, we transferred the plasmid to the BL21(DE3) expression strain to induce the expression of the AbOmpA protein by IPTG. Next, we purified the AbOmpA protein via affinity chromatography (Ni-IDA resin) (Fig S1–3). Using AbOmpA as the target, we screened the PH.D-7 PH.D-12, and PH.D-C7C peptide libraries by solid state screening. After four rounds of binding–washing– elution–amplification, six repeats were obtained, and the corresponding peptides were synthesized (Table 1). We named three peptides from the PH.D.-12 peptide library P90, P91, and P92 and three peptides from the PH.D.-C7C peptide library as P559, P560, and P609. The surface plasmon resonance results revealed that among these peptides, P92 exhibited the strongest binding force, with an equilibrium dissociation constant (KD) of 7.84 nM, followed by P90 with an equilibrium dissociation constant of 1.75 μM. P91 and P609 demonstrated moderate binding force with AbOmpA, whereas P559 and P560 displayed weaker binding force with AbOmpA (Table 1).

**TABLE 1.** Peptides obtained by phage display technology.

| Peptide | sequence | MW | Purity | RR | Avg KD (M) |
| --- | --- | --- | --- | --- | --- |
| P90 | GKWTGYGGVIHD | 1,289.41 | 97.76% | 6% | 1.75E - 06 |
| P91 | STLTSRAHTTEI | 1,316.43 | 98.03% | 6% | 2.21E - 05 |
| P92 | QMGFMTSPKHSV | 1,349.60 | 96.31% | 36% | 7.84E - 09 |
| P559 | CTPRSANYC | 1,012.16 | 97.06% | 9% | 2.11E - 03 |
| P560 | CGTNPIKKC | 961.20 | 96.89% | 12% | 1.08E - 02 |
| P609 | CDGRPDRAC | 990.11 | 95.39% | 9% | 5.43E - 06 |

MW: molecular weight; RR: repetition rate.

Next, we investigated the direct impact of KD value on the antibacterial activity of these antiAbOmpA peptides. The drug-sensitive strain, *A. baumannii* (ATCC 19606), and MDR *A. baumannii* (XJ859) were cocultured with human non-small cell lung cancer cells (A549 cells) for a duration of 2 hours. Subsequently, we examined the effects of P90, P91, and P92 on bacterial adhesion and invasiveness by quantifying the number of bacteria adhering to the surface of A549 cells as well as those entering A549 cells. As depicted in Fig lA and B, treatment with 10 μM P92 resulted in a significant reduction in the adhesion rate (approximately 75% and 90% reduction, respectively) and invasion rate (around 85% and 87% reduction, respectively) for these strains. P90 exhibited the second most significant effects, while P91 exhibited the least impact. These findings aligned with our affinity assay results, indicating that the antibacterial efficacy of the antiAbOmpA peptide was dependent on its binding strength to the target. Furthermore, growth curve analysis demonstrated that exposure to 20 μM P92 had no direct influence on the growth pattern of *A. baumannii* ATCC 19606 or MDR *A. baumannii* XJ859 (Fig 1C). Moreover, the results of Cell counting kit 8 indicated that A549 cells demonstrated an approximately 100% survival rate following treatment with experimental concentrations of P92 for a duration of 24 hours (Fig 1D). These outcomes suggest that although P92 does not possess bactericidal properties, it effectively reduces bacterial pathogenicity upon binding with OmpA.

**FIG 1.**
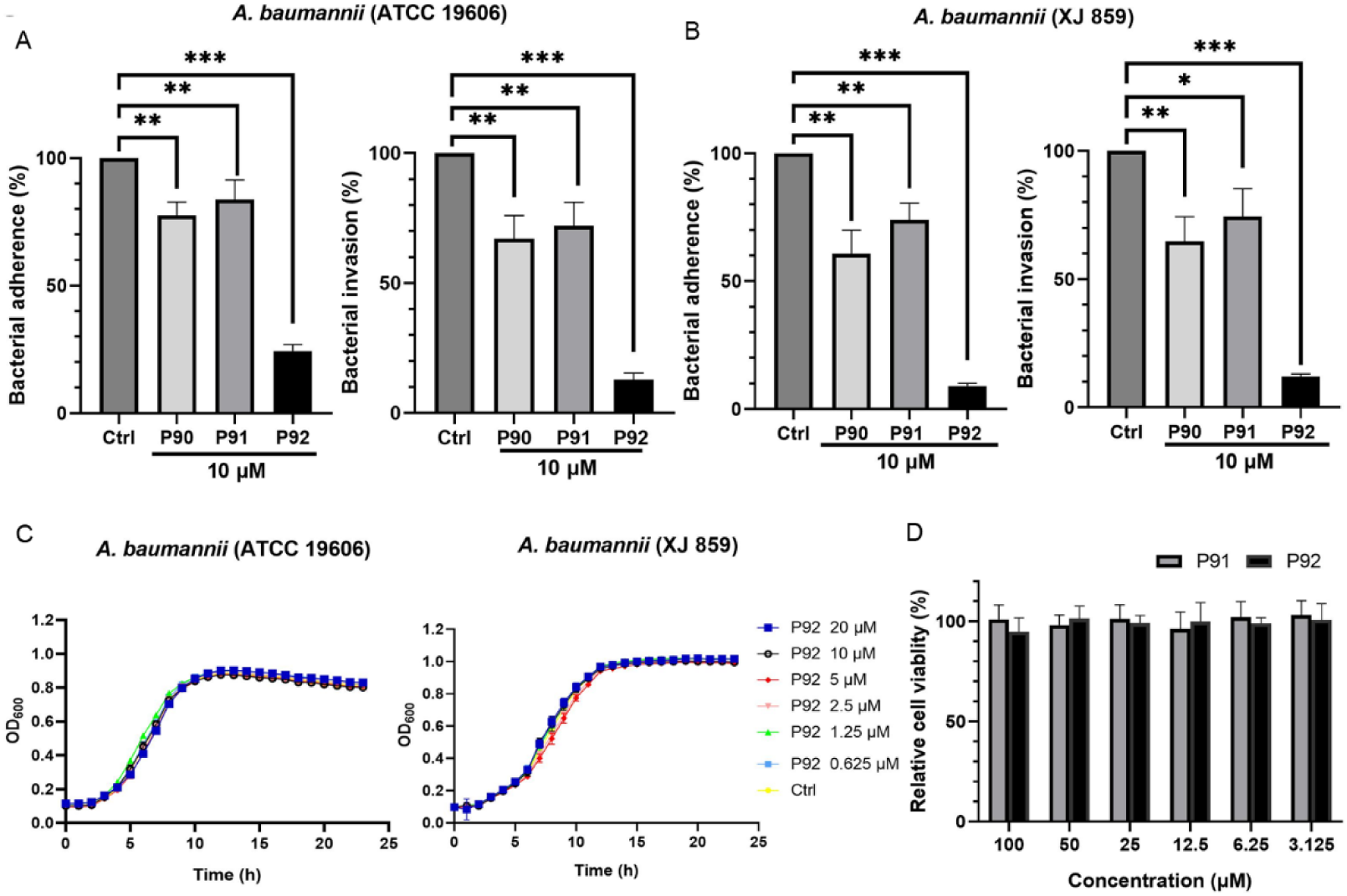
The inhibitory effect of peptides on the adhesion and invasion of *A. baumannii*. (A, B) Inhibition of P90, P91, and P92 on the invasiveness or adhesion of *A. baumannii* ATCC 19606 (A) and XJ859 (B). (C) The growth of *A. baumannii* ATCC 19606 and XJ859 treated with different concentrations of P92. (D) Cytotoxicity of P92 toward A549 cells. N = 9, * *P* < 0.05, ** *P* < 0.01, *** *P* < 0.001 vs. control.

### The anti-invasion and antiadhesion of P92 depended on OmpA

As P92 exhibited the highest affinity with AbOmpA, as well as the strongest anti-invasion and adhesion effect, we focused on the antibacterial action characteristics of P92. First, we verified that the anti-invasion and adhesion effects of P92 were concentration-dependent (Fig 2A, B). Subsequently, we compared the effects of P92 on two standard strains of *A. baumannii* and five MDR *A. baumannii* strains (Table S1). The results revealed that 10 μM P92 could sharply inhibit the adhesion and invasion of all strains except *A. baumannii* ATCC BAA-747 by more than 80%–85%, while the inhibition effect of P92 on *A. baumannii* ATCC BAA-747 was weak, with inhibition being reduced by approximately 25% (Fig 2C). The RT-PCR results further confirmed that the mRNA level of *ompA* was significantly different in *A. baumannii* from different sources, among which the mRNA level of *ompA* was the lowest in *A. baumannii* ATCC BAA-747 (Fig 2D). Correlation analysis showed that the mRNA level of *ompA* in *A. baumannii* was significantly positively correlated with the anti-invasion and antiadhesion effects of P92, and the correlation coefficients were 0.95 and 0.91, respectively (Fig 2E). These results indicated that the expression of OmpA in *A. baumannii* directly affected the effect of P92.

**FIG 2.**
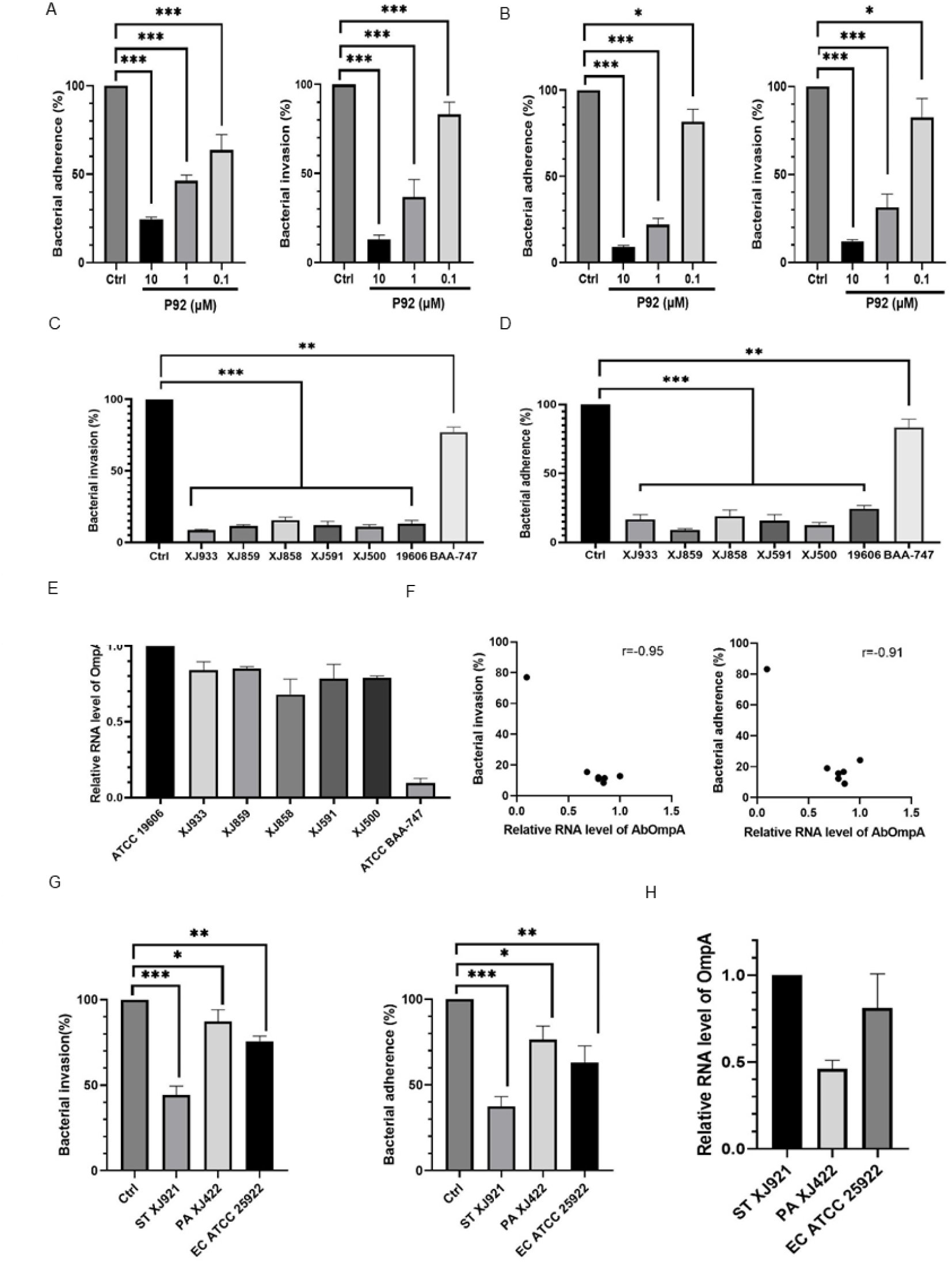
The anti-invasion and antiadhesion of P92 depends on OmpA. (A, B) The inhibitory effect of 10 μM, 1 μM, and 0.1 μM P92 on the cell invasion and adhesion of *A. baumannii* ATCC 19606 (A) and XJ859 (B). (C, D) Inhibitory effect of 10 μM P92 on the cell invasion (C) and adhesion (D) of *A. baumannii* of different strains. (E) *ompA* mRNA levels in *A. baumannii* of different strains. (F) Correlation between the inhibitory effect of P92 on the invasion or adhesion of *A. baumannii* and the level of *ompA* mRNA. (G) Inhibitory effect of 10 μM P92 on the cellular invasion and adhesion of *S. typhimurium* XJ921, *P. aeruginosa* XJ422, and *E. coli* ATCC 25922. (H) *ompA* mRNA levels in *S. typhimurium* XJ921, *P. aeruginosa* XJ422, *E. coli* ATCC 25922, and *A. baumannii* ATCC 19606. N = 9, * *P* < 0.05, ** *P* < 0.01, *** *P* < 0.001 vs. control.

Due to the widespread presence of OmpA in gram-negative bacteria and its significant role in bacterial adherence and invasion, the impact of P92 on adhesion and invasion was also investigated for three other gram-negative bacteria: *E. coli* ATCC 25922, *P. aeruginosa* XJ422, and *Salmonella typhimurium* XJ921. The findings revealed that a concentration of 10 μM P92 effectively inhibited the adhesion and invasion of various gram-negative bacteria. Notably, the strongest inhibitory effect was observed against *S. typhimurium* XJ921, with a reduction rate of approximately 40%, followed by *E. coli* ATCC 25922 and *P. aeruginosa* XJ422 (Fig 2F). Furthermore, the reverse transcription polymerase chain reaction (RT-PCR) results revealed significant variations in *ompA* mRNA levels among the three gram-negative strains, with *S. typhimurium* XJ921 exhibiting relatively high expression of *ompA*. Collectively, these findings demonstrate that P92 effectively inhibits the adhesion and invasion of other gram-negative bacteria, while its efficacy is contingent upon bacterial OmpA expression.

### Inhibitory effect of P92 on *A. baumannii* biofilm formation

Once *A. baumannii* forms a biofilm, its susceptibility to antibiotics decreases significantly, posing challenges for clinical treatment(10, 11). Considering the critical role of OmpA in the initial stage of bacterial biofilm formation, we hypothesized that P92 could act as an inhibitor of biofilm formation. Initially, *A. baumannii* ATCC 19606 and XJ859, known for their strong ability to form biofilms (Fig S4), were cultured in 96-well plates for varying durations, while the impact of P92 on biofilm formation was monitored via crystal violet staining. The results demonstrated that *A. baumannii*’s biofilms gradually matured after 48 hours of culture, with the OD_630_ reaching its peak value; however, after 72 hours of culture, the biofilms entered the exfoliation stage. Notably, compared to the control group and the group treated with 10 μM P91 or P92 for 24 hours significantly inhibited biofilm formation in the tested strains as evidenced by a reduction in the OD_630_ to <0.2 (Fig 3A). It is worth mentioning that there was no significant increase in P92’s inhibitory effect on biofilms with prolonged exposure time, which suggested that its main inhibitory action occurred during the initial stages of biofilm formation. To further validate the inhibitory effect on biofilm formation by P92, *A. baumannii* ATCC 19606 and XJ859 were subjected to different concentrations of P92 over a period of 48 hours. The outcomes revealed a concentration-dependent inhibition pattern by P92 against biofilm formation (Fig 3B). Surprisingly, even at concentrations as low as 1 μM and 0.1 μM, P92 reduced the OD_630_ levels to approximately 0.3, indicating its potent capability as an inhibitor against *A. baumannii* biofilm formation.

**FIG 3.**
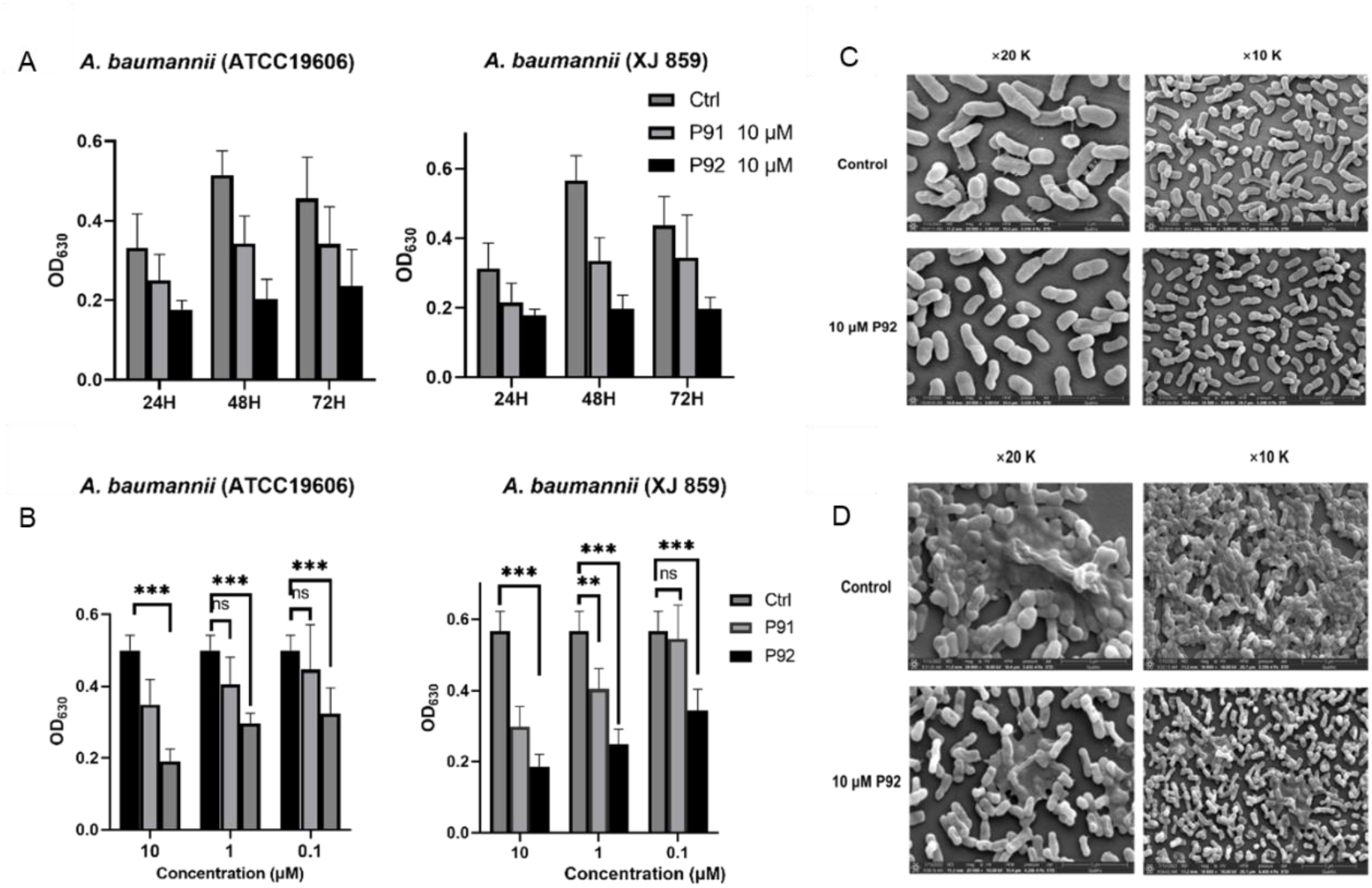
Inhibitory effect of P92 on *A. baumannii* biofilm formation. (A) Biofilm formation of *A. baumannii* ATCC 19606 or XJ859 after treatment with P92 for different times. (B) Biofilm formation of *A. baumannii* ATCC 19606 or XJ859 after treatment with 10 μM, 1 μM, or 0.1 μM P92 for 48 h. (C, D) Morphology of *A. baumannii* ATCC 19606 after treatment with 10 μM P92 for 5 h (c) or 48 h (d) investigated by SEM. Magnification: 10 k or 20 k. N = 10, ** *P* < 0.01, *** *P* < 0.001 vs. control.

To further investigate the morphological effects of P92 on *A. baumannii*, *A. baumannii* ATCC 19606 was treated with a concentration of 10 μM P92 for 5 hours. The scanning electron microscopy (SEM) results revealed that, compared to the control group, the bacterial surface in the P92 treatment group exhibited significantly reduced filiform secretions and adhesion (Fig 3C). This indicated that while P92 did not visibly affect the morphology and size of planktonic cells, it notably diminished bacterial interactions. Subsequently, slides were introduced into the culture system containing *A. baumannii* ATCC 19606 and incubated for 48 hours. SEM analysis demonstrated that, in comparison to the control group, a substantial number of bacteria on the slides within the P92 treatment group displayed dispersed distribution characteristics accompanied by a sharp reduction in biofilm formation area as well as thickness and density (Fig 3D). These findings suggest that through inhibiting bacterial adhesion, P92 may effectively impede *A. baumannii* biofilm formation.

### Protective effect of P92 on *A. baumannii*-infected cell and animal models

To confirm the overall therapeutic efficacy of P92 against infection, we investigated its protective effect on A549 cells infected with *A. baumannii*. Following a 24-hour infection period with three different strains of *A. baumannii*, the cell viability in the infected group decreased to approximately 10% compared to the control group. However, treatment with P92 and P91 significantly improved cell viability in all three strains of *A. baumannii*-infected cells. Notably, the highest level of protection was observed in the 20 μM P92 treatment group. The cytoprotective effect of P92 varied among different strains of infection; this was consistent with our findings on the anti-invasion and antiadhesion properties. Specifically, following treatment with 20 μM P92, the cell viabilities of the *A. baumannii* ATCC 19606, XJ859, and ATCC BAA-747-infected groups were 82%, 79.9%, and 42%, respectively (Fig 4A). These results further support that the protective effect of P92 on infected cells is associated with AbOmpA expression levels in bacteria.

**FIG 4.**
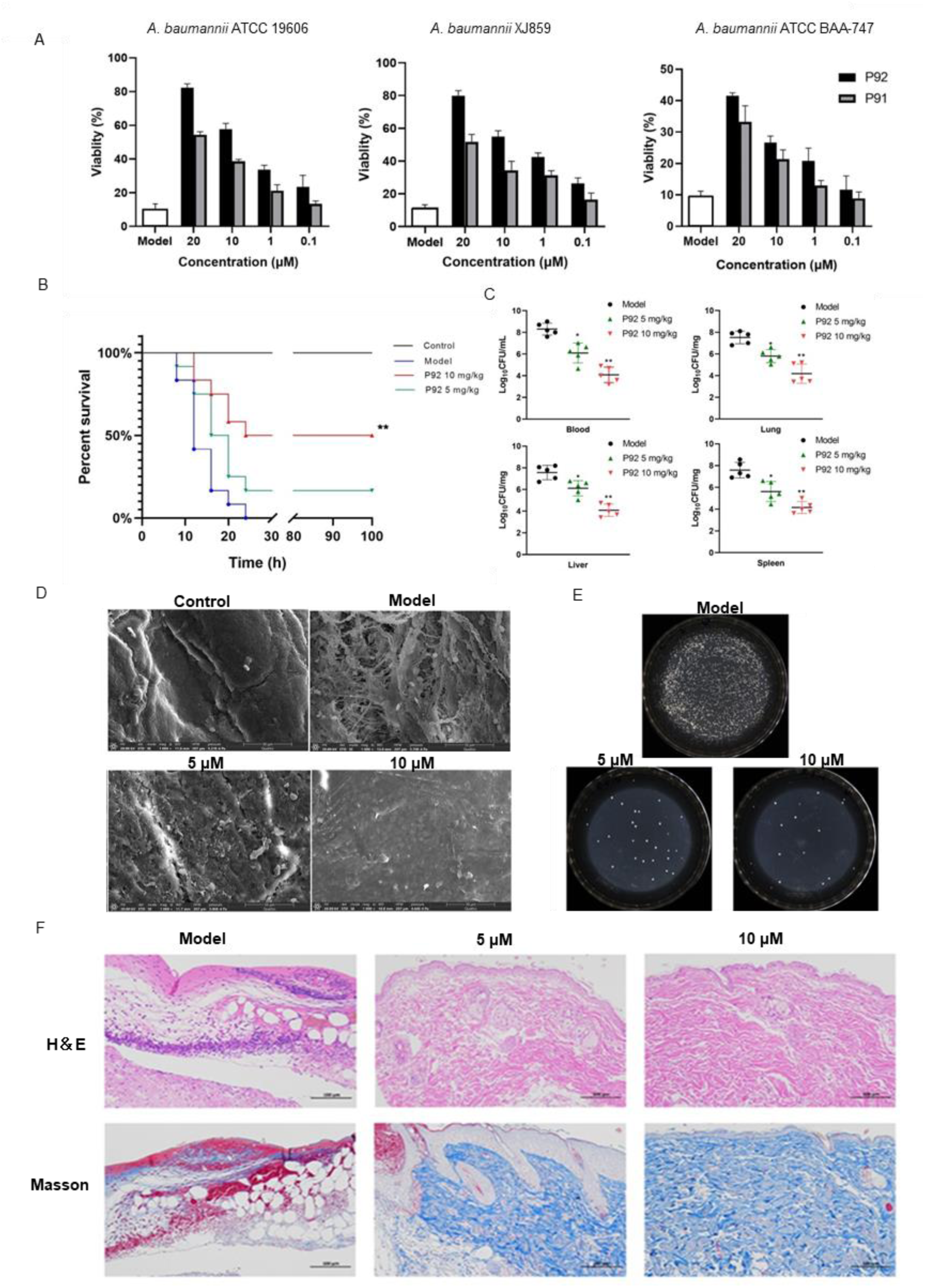
Protective effect of P92 on *A. baumannii-*infected A549 cells and animal model. (A) Viability of A549 cells infected by *A. baumannii* ATCC 19606, XJ859, or ATCC BAA-747 after treatment with different concentrations of P91 and P92; (B) Survival rate of septic mice induced by *A. baumannii* ATCC 19606 after treatment with 5 mg/kg or 10 mg/kg P92 (N = 12 per group). \*\**P* < 0.01 vs. model (log-rank test). (C) Colonization of *A. baumannii* inocula in blood, lung, liver, and spleen of infected mice (N = 5 per group), \**P* < 0.05 vs. model. \*\**P* < 0.01 vs. model. (D) Morphological observation of mouse skin wound biofilm (*A. baumannii* ATCC 19606) of control group (uninfected), model group, 5 μM, and 10 μM P92 group by SEM. (E) Colony counting of mice wound exudate of model group, 5 μM, and 10 μM P92 group. (F) HE staining and Masson three-color staining of mouse model group, 5 μM, or 10 μM P92 group. Scale: 500 μm.

Subsequently, a mouse model of sepsis was established by intraperitoneally injecting 100 µL of bacterial suspension containing *A. baumannii* ATCC 19606 at a concentration of 5 × 10^8^ CFU/mL. Then, the mice were treated with varying doses of P92 at 1 h, 7 h, and 13 h postinfection. Consequently, all mice in the control group succumbed within 24 h, whereas the survival rates increased to 16.7% and 50% for mice treated with P92 at doses of 5 mg/kg and 10 mg/kg, respectively (Fig 4B). Furthermore, P92 exhibited a dose-dependent reduction in CFU count in the blood and organs of sepsis-infected mice. Specifically, the model group displayed approximately a 2.0 × 10^8^ CFU/mL bacterial count in the blood, while the lung, liver, and spleen had counts of around 4.0 × 10^7^ CFU/mg each. Following treatment with P92 at a dosage of 10 mg/kg, the bacterial counts decreased to approximately 1.3–2.2 × 10^4^ CFU/mg across the blood, lung, liver, and spleen (Fig 4C). These findings unequivocally demonstrate that even severe sepsis-infected mice can benefit significantly from P92 administration.

Furthermore, a skin wound infection model of *A. baumannii* ATCC 19606 was established to assess the protective effect of P92. The SEM images revealed that, compared to uninfected skin wounds, the model group exhibited evident filamentous membrane-like structures and a large number of bulbous bacteria 48 h after infection (Fig S5). This indicated the secretion of abundant polysaccharides and other substrates by the bacteria at the wound site, resulting in biofilm formation. Immediately following infection, different concentrations of P92 solution were applied three times a day to mouse skin wounds. After 2 days of continuous treatment, wound exudates were collected for bacterial colony counting. The results confirmed a significant reduction in bacterial count in the administration group compared to the control group, demonstrating concentration-dependent effects of P92 (Fig 4D). The SEM images revealed noticeable filamentous membrane-like structures on the wound surface in the control group; however, these structures as well as the bacterial count were significantly reduced in the low-dose P92 administration group, with partially flattened wounds observed. Furthermore, almost completely flat wound surfaces were observed in the high-dose P92 administration group (Fig 4D). These findings suggest that P92 effectively reduces bacterial load in skin infections and inhibits biofilm formation (Fig 4E). Moreover, tissue analysis performed two days following administration of P92 indicated early wound healing progress. Hematoxylin and eosin (HE) staining demonstrated reduced inflammatory cell infiltration and alleviated inflammation in the administration group compared to the controls. Masson staining revealed an increased blue fiber area and significantly higher collagen deposition in the P92-administered groups than in controls (Fig 4F). These data indicate that P92 reduces biofilm formation and promotes early wound healing by inhibiting bacterial colonization of skin wounds.

## Discussion

Bacterial resistance to antibiotics is inevitable because of the direct impact of traditional antibiotics on essential molecular functions that are required for bacterial survival. For example, penicillin G inhibits the activity of transeptidase by binding with penicillin-binding proteins, thus obstructing cell wall synthesis and inducing tremendous survival pressure in bacteria. Consequently, bacteria are constantly driven to generate mutations to adapt to environmental changes, and it is not an exaggeration to assert that drug resistance is an unavoidable consequence of bacterial survival. However, the era of abundant antibiotic discovery has long passed; as we enter the 21^st^ century, finding new antibiotics becomes increasingly challenging. Therefore, it is unrealistic to address the issue of bacterial resistance solely through the continuous pursuit of novel antibiotics(12–14). Particularly concerning the infiltration barrier and efflux mechanism of gram-negative bacteria, MDR gram-negative bacteria continue to emerge as a major cause of mortality among ICU patients(15). While small molecules th at inhibit or kill bacteria have been a mainstay of antimicrobial therapy in the past, “unconventional” approaches are now gaining momentum as alternative strategies for overcoming scientific obstacles posed by traditional antibiotics against gram-negative bacteria. These unconventional approaches include antivirulence strategies, phage therapies, microbiome modification therapies, antimicrobial vaccines (16, 17). Among them, the antivirulence strategy aims to play a distinctive role by interfering with toxin production and virulence factor secretion by bacteria, preventing bacteria from adhering to host cells and forming biofilms, interrupting or inhibiting bacterial communication, or downregulating virulence(18, 19). Currently, there are dozens of related research projects at various stages ranging from preclinical studies to clinical trials(20, 21).

The outer membrane proteins (OMPs), serving as a critical virulence factor in *A. baumannii*, belong to a distinctive class of integral membrane proteins that are firmly anchored in the OM and possess a ß-barrel structure composed of 8 to 26 chains(22). Among the OMPs of *A. baumannii*, OmpA has been extensively studied as a virulence factor that plays a critical role in regulating adhesion, invasiveness, and biofilm formation in *A. baumannii*, as well as the host’s immune response(4). Studies on drugs targeting AbOmpA have been reported for a long time. For example, AOA-2 is a cyclic hexapeptide obtained via virtual screening(9, 23); however, its affinity with the target is far lower than that of P92 obtained through phage screening, whose KD value is 7.84 nM. We discovered that P92 inhibits invasion and adhesion in various gram-negative bacteria with varying degrees of effectiveness. This discrepancy arises from the fact that the OmpA family consists of high copy number membrane porins found in the OMPs of gram-negative bacteria. While its amino acid sequence remains highly conserved (>89%) among different clinical isolates of *A. baumannii*, it differs across other gram-negative bacteria. Consequently, P92 specifically targets AbOmpA and exhibits strong inhibitory effects on *A. baumannii* while displaying weaker effects on other gram-negative bacteria such as *P. aeruginosa* and *E. coli.* Additionally, we observed significantly reduced efficacy of P92 against *A. baumannii* ATCC BAA-747 because of its low OmpA copy number. However, it should be noted that all MDR clinical isolates used in this study had high OmpA copy numbers, indicating that AbOmpA is an ideal target against *A. baumannii*.

We evaluated the overall therapeutic efficacy of P92 against infection in a mouse skin infection model and a mouse sepsis model. Notably, in mice with severe sepsis, P92 did not confer complete survival benefits, even at high doses (10 mg/kg), as the survival rate was only 50%; this result specifically distinguishes antivirulence strategies from traditional antibiotics because the former does not directly kill or inhibit bacterial growth. Consequently, they are not intended as an “alternative” to antibiotics but are designed as an adjunct therapy alongside antibiotic treatment. Furthermore, we attempted combination therapy using P92 and antibiotics for sepsis treatment; however, unfortunately, no efficacy of P92 was observed based on the available results (results not shown). This highlights that experimental design and selection of observation indicators are pivotal for successful implementation of antivirulence strategies as adjunct treatments for infection. Nevertheless, we cannot disregard the effectiveness of P92, as a virulence factor inhibitor, for the treatment of septic infections because of its dose-dependent reduction in bacterial load within the blood and organs along with significant improvement in sepsis-affected mouse survival rates. Moreover, P92 appears more suitable for topical application against skin bacteria rather than serving as an adjunct therapy for systemic antibacterial infections. P92 can reduce bacterial burden without inducing drug resistance. Moreover, it significantly inhibits biofilm formation in skin wounds, which greatly facilitates early wound healing. In conclusion, these findings strongly suggest that P92 acts as a highly effective OmpA inhibitor leading to substantial attenuation of bacterial virulence and pathogenicity, while representing a promising novel antibacterial molecule.

However, it is important to acknowledge that virulence factors such as AbOmpA exhibit species or even strain specificity and demonstrate variable conformity within and between bacterial species. Furthermore, the expression of virulence genes may be influenced by environmental conditions or the site of infection, as well as the temporal progression of pathophysiological processes. The intricate biodiversity presents a significant challenge in translating initial findings into clinical scenarios. d

In conclusion, our results demonstrated remarkable therapeutic effects of anti-AbOmpA peptide P92 against various models of *A.baumannii* infection without causing significant harm to bacterial or mammalian cell viability even at high concentrations; dffective inhibitor of virulence factors with promising prospects for anti-infection therapy. However, given the limitations of targeted bacterial factor strategies, it is imperative to conduct further evaluations on the efficacy of P92 against a broader range of *A. baumannii* sources. Additionally, investigating the combined effects of P92 and antibiotics on bacterial resistance and anti-infective efficacy is crucial in comprehensively assessing the reliability of antimicrobial virulence factor-based therapy.

## MATERIALS AND METHODS

### AbOmpA synthesis

The AbOmpA protein was synthesized by Minyan Biotechnology Co. In brief, the amino acid sequence of the AbOmpA protein was optimized using MaxCodonTM Optimization Program (V13), a newly developed codon optimization software by MINGYAN BIOLOGICAL. Subsequently, the *ompA* gene was inserted into the expression vector pET30a through whole gene synthesis and restriction enzyme cleavage sites Ndel and HindIII. The accuracy of the final expression vector was confirmed through digestion and sequencing analysis. Following that, the AbOmpA protein was transferred to BL21(DE3) expression strain and induced with IPTG for expression purposes. Finally, purification of the AbOmpA protein was achieved using affinity chromatography with NiIDA resin.

### Peptides synthesis

The anti-AbOmpA peptides were synthesized by Qiangyao Company. In brief, the peptides were synthesized in a sequential manner from the C-terminal to the N-terminal of the sequence. Subsequently, crude products underwent purification to achieve the required purity using high-performance liquid chromatography (HPLC). Finally, after purification, they were lyophilized and concentrated into a white powder.

### Biopanning of phage-displayed peptides

The direct target coating method was employed to screen Ph.D.-7 peptide library, Ph.D.-12 peptide library, and Ph.D.-C7 peptide library (New England Biolabs, Ipswich, MA, USA) for the identification of AbOmpA binding phage. In the initial round of biological screening, the 96-well plate was coated with AbOmpA protein and incubated overnight. Subsequently, after blocking with a blocking buffer (0.1 M NaHCO_3_, 5 mg/mL BSA), it was incubated with the phage library at a final concentration of 1.5 × 10^11^ (100 uL/well) at room temperature for 0.5 h. The unbound bacteriophages were then washed ten times with TBST (TBS containing 0.1% Tween-20). Following this, an elution buffer (100 μg/mL AbOmpA TBS solution) was added and allowed to elute for one hour at room temperature. The eluted bacteriophages were subsequently titrated and amplified for the second round of biological screening. After four rounds of biological screening, the phage storage solution was prepared using the eluted phage system, and the phage genomic DNA was isolated and sent to Orco Biotechnology Co., Ltd for nucleotide sequencing.

### SPR test

The AbOmpA protein solution was subjected to dialysis with PBS in order to remove detergents, glycerol, and other additives. After concentration, the protein concentration was determined using a BCA protein quantitative kit. The AbOmpA protein was then diluted with PBS to achieve the desired printing concentration and printed onto a 3D optical cross-linking chip (AD1520 chip array printer). Each sample was printed with four repetitive points, while rapamycin served as the positive control point. Five concentration gradients of six types of peptides were diluted with PBST to concentrations ranging from 10 nM to 2560 nM. All samples were tested on the chip surface at a flow rate of 0.5 μ L/s from low to high concentration under conditions where reaction temperature was set at 4 C for binding time of 10 min followed by dissociation time of 6 min, and then the Glycine-HCl solution of pH2.0 was used as regeneration solution at a flow rate of 2 μ L / s.

### Susceptibility test

The MICs of imipenem, ciprofloxacin, ceftazidime, gentamicin, ampicillin, and Polymyxin B for both standard and clinical isolates of *A. baumannii* strains were determined using the microdilution assay in accordance with CLSI guidelines(24).

### Growth curve

*A. baumannii* ATCC 19606 and *A. baumannii* XJ859 were cultured to logarithmic growth stage, and then diluted 100-fold with MHB broth to a concentration of about 1×10^6^ CFU/mL. 150 μL of multiply diluted peptide was added to each well of a well plate for the growth calibrator at the concentration of 1.25 μM∼40 μM,. Then 150 μL of bacterial suspension was added to each well, and the concentration gradient of peptide was 0.625 μM∼20 μM. The OD_600_ value of each well was measured by the growth calibrator at different time points, and the temperature was set at 37 ℃, and the time interval was 1 h, and it was measured continuously for 23 h.

### Cell culture

The A549 cell line was obtained from our laboratory. Cells were grown in DMEM supplemented with FBS (10%) at 37 °C in a humidified 5% CO_2_ incubator.

### Cellular viability

#### Toxicity to A549 cells

A549 cells were seeded at a density of 5000 cells/well in 96-well plates and incubated overnight at 37 ℃ with 5% CO_2_. The supernatant was removed, and P92 and P91 solutions (100 μL) were added to each well, which had been diluted with DMEM medium to concentrations ranging from 3.125 μM-100 μM. After incubation for an additional 24 h, CCK-8 solution (10 μl; Dojindo, Kumamoto, Japan) was added according to the manufacturer’s instructions. Absorbance was measured at 450 nm after continued incubation for another two hours at 37 °C.

#### Protective effect on infected cells

A549 cells were infected with 5^10^6^ CFU/mL of *A. baumannii* ATCC 19606, *A. baumannii* ATCC BAA-747 and *A. baumannii* XJ859 strains pretreated with P92 (0.1, 1, 10 and 20 μM, 1 h) for 24 h with 5% CO_2_ at 37 °C. Before assessing bacterial cytotoxicity, viable bacteria were removed from the A549 cell cultures by washing the cells five times with pre-warmed PBS. Subsequently, cellular viability was evaluated as described above.

### Adhesion assays

*A. baumannii* strains were incubated with peptides (10 μM, 1h), and subsequently added to A549 cells for a 2-hour incubation at 5% CO_2_ and 37 °C. Following this, the infected A549 cells were washed five times with pre-warmed PBS and lysed using 0.3% Triton X-100. Diluted lysates were plated onto MHA and incubated at 37 °C for 24 hours to enumerate developed colonies and determine the number of bacteria that adhered to A549 cells.

### Invasion assays

*A. baumannii* strains were incubated with peptides (10 μM, 1h) and subsequently added to A549 cells for a 2-hour incubation at 5% CO_2_ and 37 °C. Following this, gentamicin was introduced at a final concentration of 200 ug/mL and treated for 30 minutes. The infected A549 cells were then washed five times with pre-warmed PBS and lysed using 0.3% Triton X-100. Diluted lysates were plated onto MHA agar plates and incubated at 37 °C for a period of 24 hours to enumerate the developed colonies, thereby determining the number of bacteria that adhered to the A549 cells.

### Biofilm formation

We used an overnight culture of 2 reference strains and 5 clinical isolates of *A. baumannii,* cultured in fresh Luria Bertani (LB) in 96 well plates that were incubated in presence or not of 10 μM P92 without shaking at 37 °C during 24 h, 48 h and 72 h. Biofilm was stained with crystal violet 1% (v/v) and quantified at 630nm after solubilization with ethanol 95%.

### Scanning electron microscope (SEM)

An overnight culture of *A. baumannii* ATCC 19606 was centrifuged at 5000 rpm for 10 min, wash twice in sterile PBS buffer, resuspend the treatment in PBS. Sterile cell crawlers were placed in 24-well plates, and each well was added with a bacterial solution of *A. baumannii* ATCC 19606 (final concentration of 5×10^6^ CFU/ mL) and a P92 solution (final concentration of 10 μM). After incubating for 48 hours, pre-fixation was performed by adding 1 mL of a 3% glutaraldehyde solution and leaving it overnight at 4°C. In addition, two tubes were taken where the same bacterial and P92 solutions were added and incubated for four hours. The samples were then centrifuged at 12,000 rpm for ten minutes to discard the supernatant before adding the glutaraldehyde solution (1 mL) followed by an overnight stay at 4°C. Finally, these samples were were further processed and then examined under scanning electron microscopy

### Real-time RT–PCR studies

The 6 standard and clinical *A. baumannii* strains, *E. coli*, *P. aeruginosa* and *S. typhimurium* were analysed for the mRNA level of the o*mpA* gene. DNase-treated bacterial RNA was isolated following the protocol of the M5 RNA Extraction Kit (Mei5 Biotechnology Co., Ltd). Real-time RT–PCR was performed with a ChamQSYBR qPCR Master Mix Real-Time PCR Detection System (Novozymes Biotechnology Co.). The primers and probes given in Table S2 were used in this study.

### Animals

The BALB/c male mice (18–22 g) were obtained from the Animal Experimentation Centre of the Air Force Military Medical University, and the animal experiments of this study complied with the regulations of the Ethics Committee of the Air Force Military Medical University and were approved.

### Sepsis model

A murine peritoneal sepsis model was established by intraperitoneal inoculation of mice with *A. baumannii* ATCC 19606 strain and treatment with P92 monotherapy. Briefly, animals were infected intraperitoneally with 0.1 mL of 5×10^8^cfu/mL of *A. baumannii* ATCC 19606. Fifty-five mice were randomly ascribed to the following groups: model group (without treatment after infection) (17 mice); 10 mg/kg or 5 mg/kg P92 administered intraperitoneally at 1 h, 7 h and 13 h after bacterial inoculation (17 mice per group); control group (4 mice). Mortality was recorded over 100 h. Five mice were taken from each group except control group at 18 h, blood samples were obtained by by removing the eyeball. The aseptic thoracotomies were performed, spleen, liver and lungs were aseptically removed and homogenized in 1 mL of sterile PBS solution. Ten-fold dilutions of the blood and homogenized spleen, liver, lungs were plated onto MHA for quantitative cultures (log cfu/mg of spleen, liver or lung, log cfu/mL of blood).

### Skin infection model

Balb/c mice were anesthetized with intraperitoneal injection of 3.5% chloral hydrate at a dosage of 10 μL/g body weight, Subsequently, they were positioned in a prone position on the fixed table and their back hair was shaved off. The skin was then disinfected and a hole with an 8mm skin punch was created at the center of the back, followed by removal of the entire layer of skin. The concentration of *A.baumannii* ATCC19606 was adjusted to 10°CFU/mL. After weighing, Balb/c mice were randomly divided into three groups: a negative control group where 30 μL of bacterial solution was applied to the mouse wound and 30 μL of PBS was immediately smeared after the bacteria solution dried; in the low and high dose group, 30 μL of bacterial solution was applied to the mouse wound followed by immediate application of 5 μM or 10 μM P92 in 30 μL. The medication administration occurred three times daily and observations were made after two days. Subsequently, SEM analysis was conducted on the skin surface. The wound and surrounding skin were excised, fixed in a 4% paraformaldehyde solution, and part of the skin tissue underwent SEM observation for biofilm growth. Another portion of the skin tissue was embedded in paraffin for HE and Masson staining analysis. The scab on each mouse’s back was lifted and tissue exudate samples were collected using sterile cotton swabs approximately ten times per sample. These swabs were soaked for five minutes in 1 mL PBS before being repeatedly squeezed to extract any remaining fluid which would then be transferred onto sterile MHA plates using a volume of one hundred microliters (uL). The plate contents would be evenly spread using a spreader before incubating at a temperature of thirty-seven degrees Celsius for eighteen hours following which colony counts would be performed.

## Acknowledgments

This research was supported by the National Natural Science Foundation of China (Grant No. 82273978) and Shaanxi Province key research and development plan (2023-YBSF-629).

## Supplementary Information

**FIG S1.**
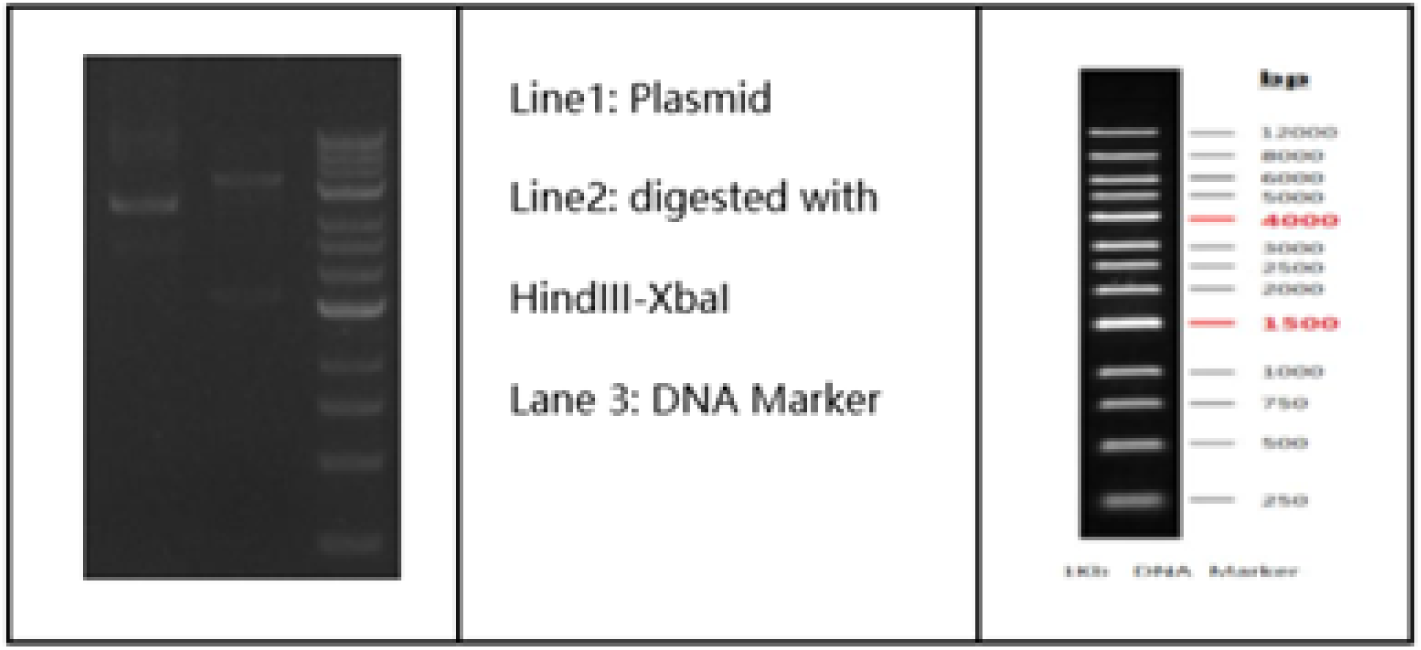
Results of plasmid quality control.

**FIG S2.**
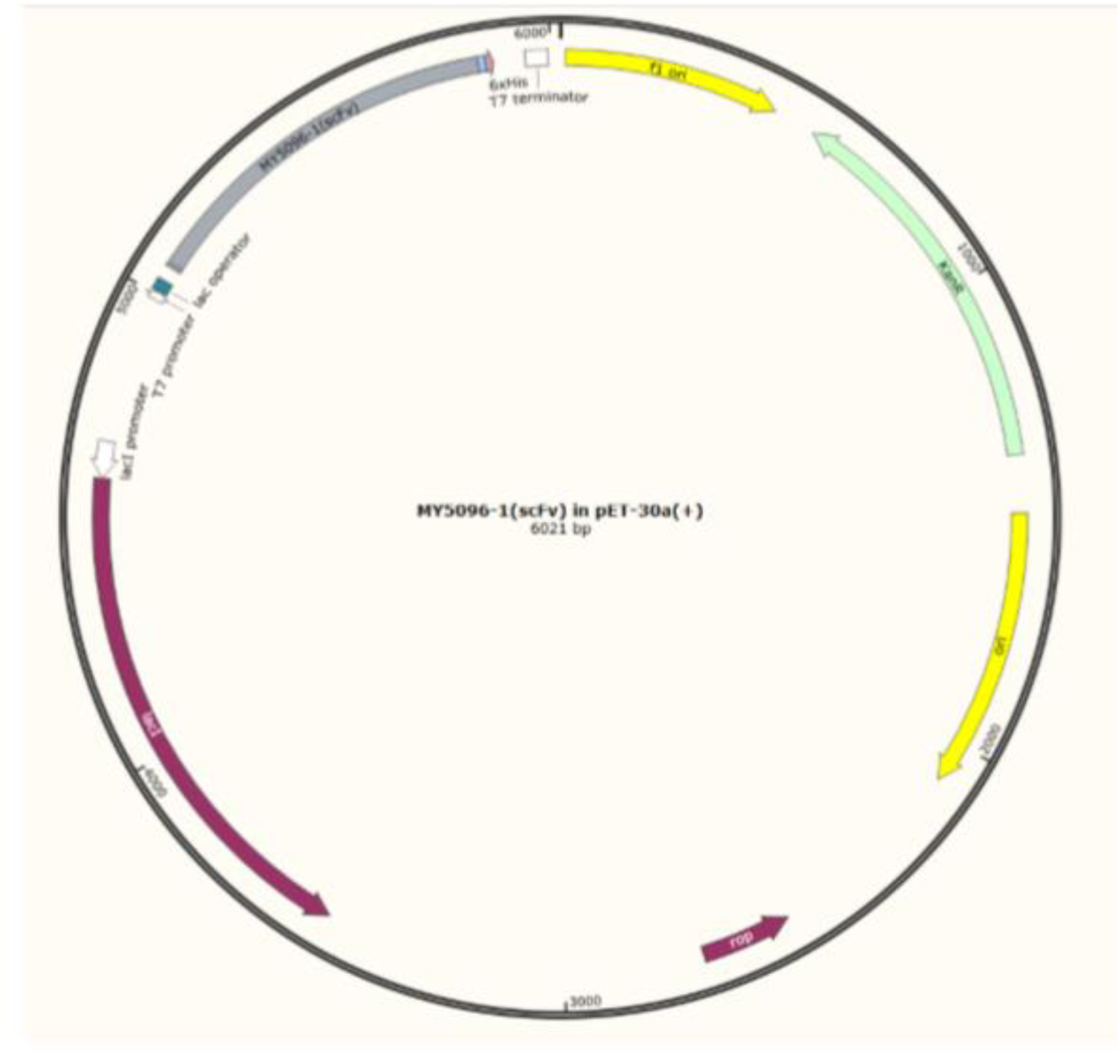
Plasmid structure.

**FIG S3.**
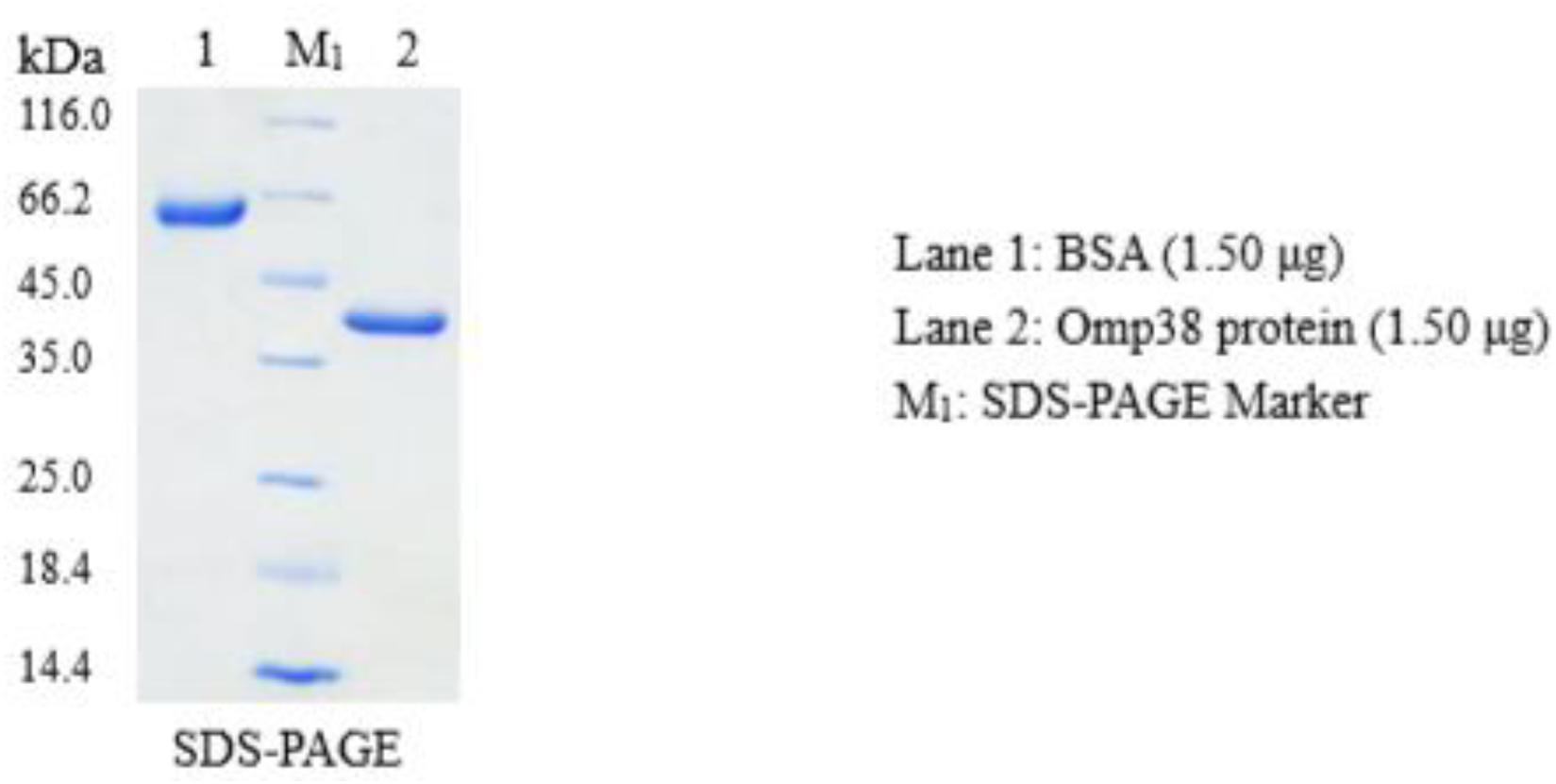
Results of AbOmpA quality control.

**FIG S4.**
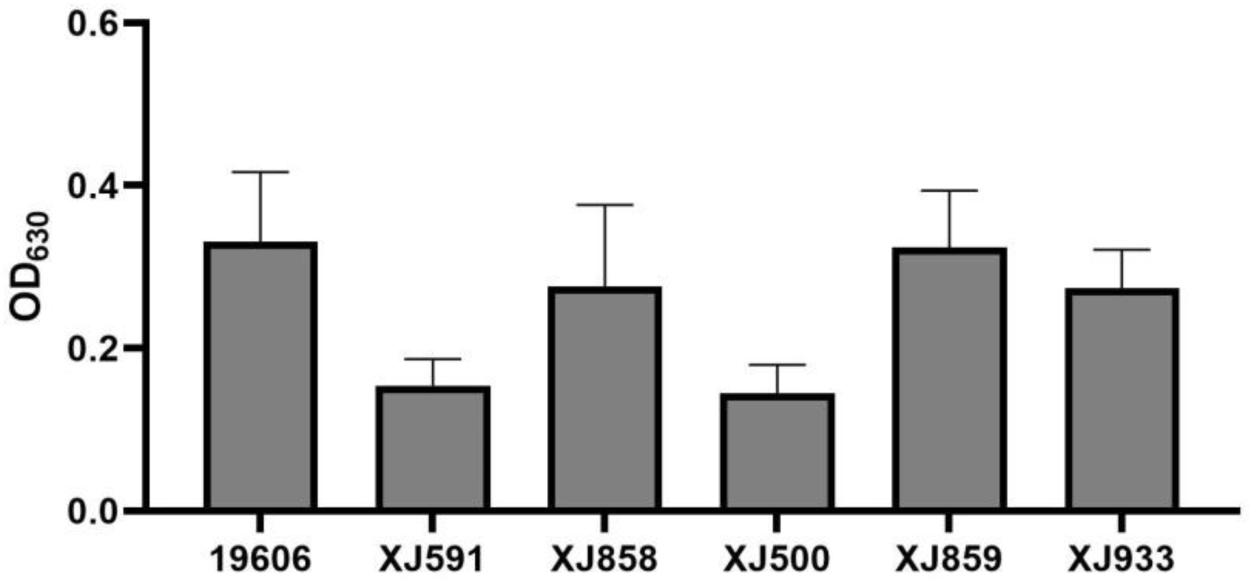
Biofilm-forming capacity of *A.baumannii*.

**FIG S5.**
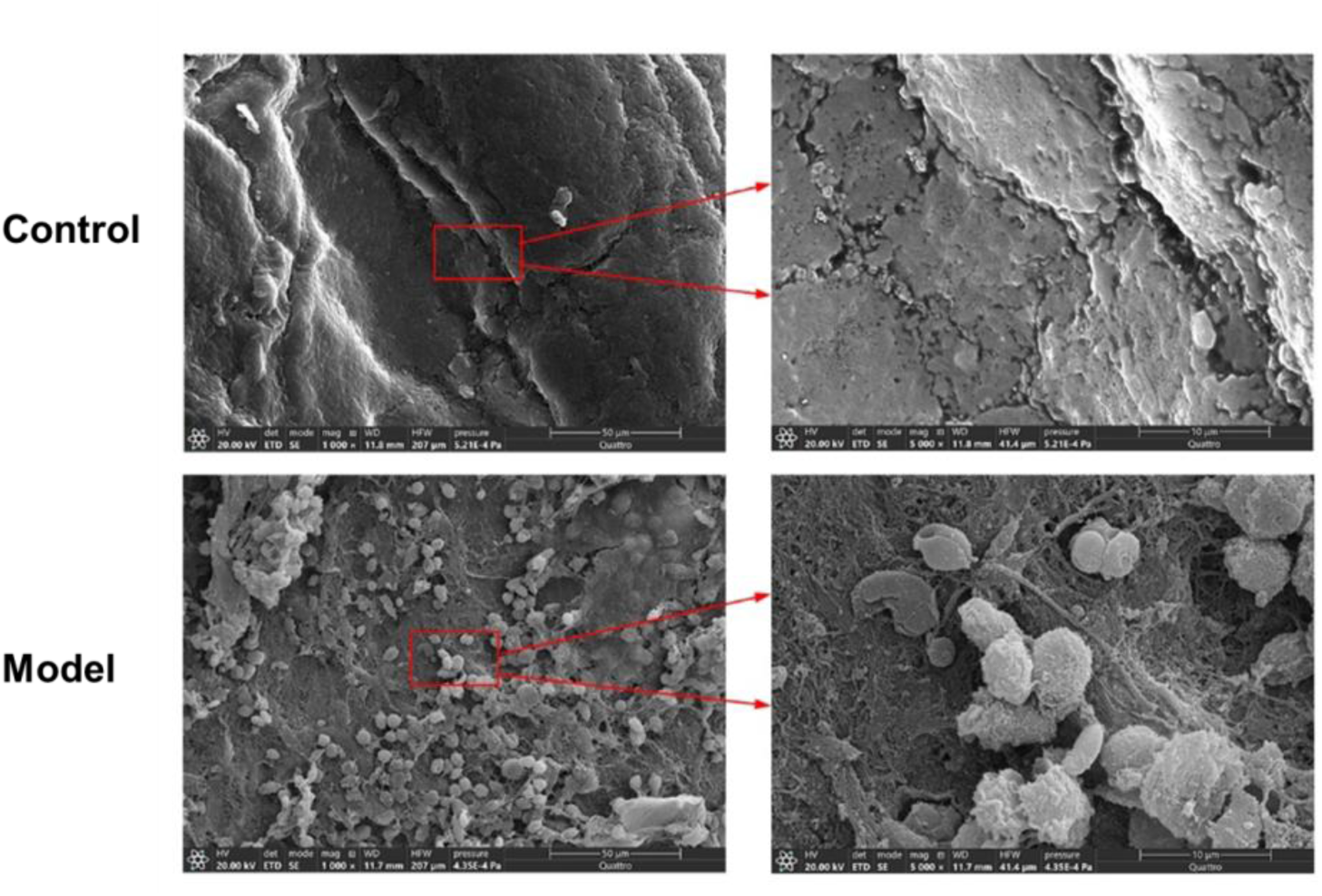
Morphological identification of a mouse skin wound biofilm infection model by scanning electron microscopy. The wound surface of the normal control group was smooth (top left and top right), and there were a lot of bacteria and membrane-like structures in the model group (bottom left and bottom right).

**TABLE S1.** MICs of antibiotics against clinial isolates of *A. baumannii*.

| Antibiotics<br>MIC($\mu\text{g/mL}$) | XJ933 | XJ859 | XJ858 | XJ591 | XJ500 | ATCC<br>19606 | ATCC<br>BAA-747 |
| --- | --- | --- | --- | --- | --- | --- | --- |
| Ampicillin | >256 | >256 | >256 | >256 | >256 | 64 | 8 |
| Ceftazidime | 128 | 128 | 128 | >256 | >256 | 16 | 8 |
| Gentamicin | 128 | 64 | 4 | >256 | 128 | 8 | 0.5 |
| Imipenem | 64 | 64 | 4 | 32 | 64 | 0.25 | 0.25 |
| Polymyxin B | 16 | 16 | 8 | 16 | 0.5 | 0.5 | 2 |
| Ciprofloxacin | 16 | 32 | 4 | 32 | 32 | 1 | 0.25 |

**TABLE S2.**
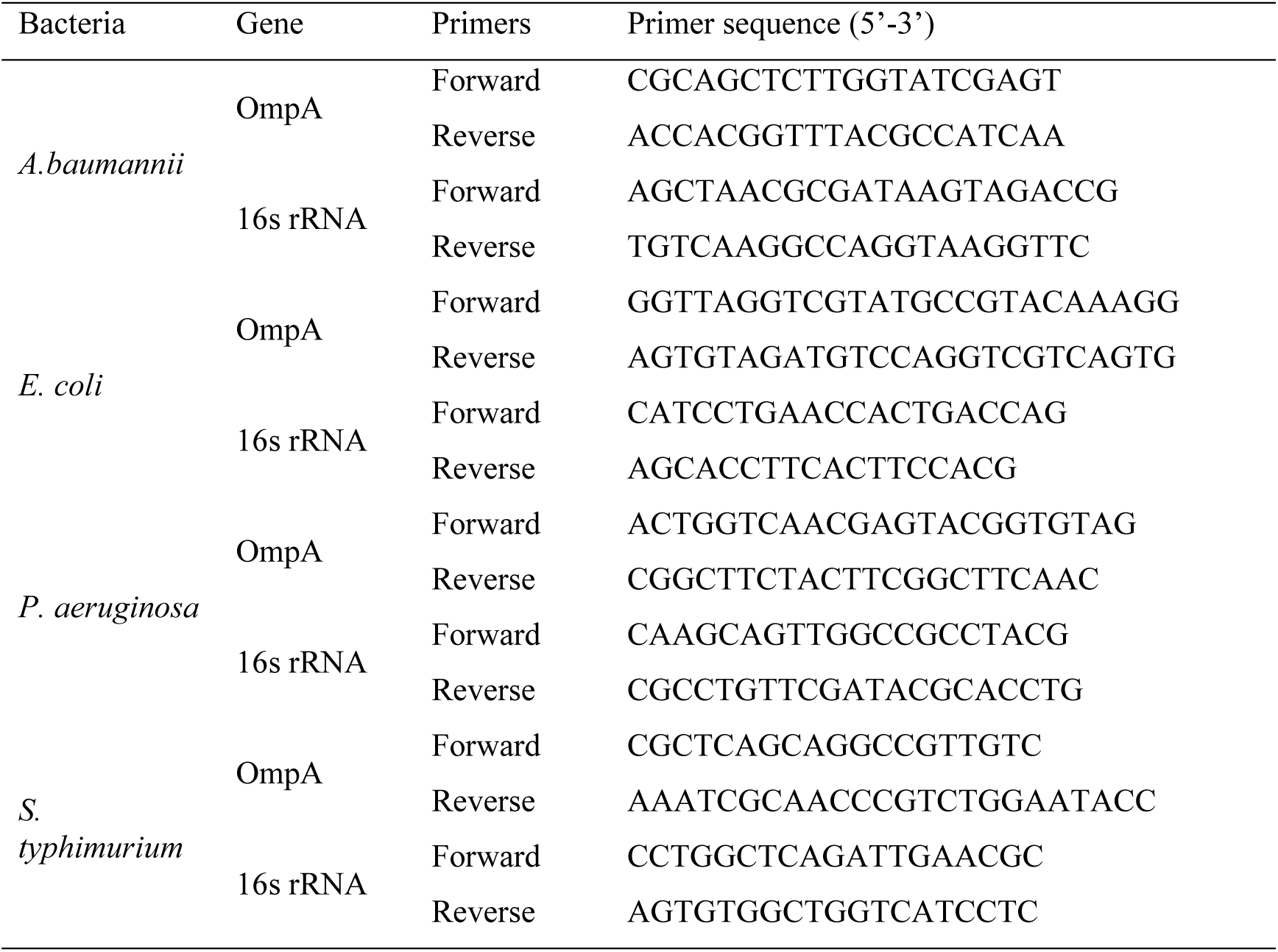
Primers sequences used for real-time PCR.

